# A low concentration of a sustainably obtained blueberry extract improves the post-thawing motility of cryopreserved bull spermatozoa

**DOI:** 10.64898/2026.03.31.715696

**Authors:** Gabriela García Blanco, Cristina Fra-Hernánez, Joana F. do-Vale-Rabaça, Laura Pariente-Martín, Marta Veza-Cuenca, Estela Fernández-Alegre, Beatriz Martín-Fernández, J. Néstor Caamaño, J. Ramiro González-Montaña, Marta Lores, Felipe Martínez-Pastor

**Affiliations:** INDEGSAL, Universidad de León, León, Spain; Dept. of Molecular Biology (Cell Biology), Universidad de León, León, Spain; IMAPOR, Universidad de León, León, Spain; Bianor Biotech SL, León, Spain; Department of Medicine, Surgery and Veterinary Anatomy, Universidad de León, Spain; Selección y Reproducción Animal-SERIDA, Principado de Asturias, Gijón, Spain; LIDSA, Department of Analytical Chemistry, Nutrition and Food Science, Faculty of Chemistry, Universidade de Santiago de Compostela, Santiago de Compostela, Spain

**Keywords:** bull, cryopreservation, sperm motility, blueberry, natural extracts, antioxidants

## Abstract

Natural extracts could improve sperm storage and artificial insemination (AI). This study, for the first time, evaluates the suitability of a blueberry extract (*Vaccinium corymbosum*) obtained from pomace using a sustainable methodology as a supplement for bull semen extenders. Cryopreserved semen doses from eight bulls were combined in 9 pools (3 bulls/pool), supplemented with 0%, 1%, 5%, or 10% extract, and incubated up to 5 h at 38 °C. Motility was assessed hourly using OpenCASA, and the effects of treatment and time were evaluated using linear mixed-effects models. Motility was significantly better preserved with 1% extract (total and progressive motility, improved linear velocity and linearities, and decreased BCF and fractal dimension, related to hyperactivation). The effect of 5% was overall positive, but it was below 1%, whereas 10% mostly showed a negative effect. These results show that this natural extract could safely supplement bull semen extenders at least between 1% to 5%, and even help improve sperm motility. Therefore, this extract offers an opportunity to enhance cattle semen extenders using a sustainable approach, potentially improving reproductive outcomes.

## 1 Introduction

Interest in natural extracts in semen extenders has increased notably (Ros-Santaella & Pintus, 2021; Schulze et al., 2020). These extracts are rich in antioxidant and antimicrobial compounds such as polyphenols. Although their composition is complex, this is also why they exhibit beneficial properties (Gato et al., 2021). These products are especially attractive for sperm cryopreservation, a stressful process in which excessive production of oxidative species and the redox imbalance decrease sperm fertility. Moreover, antimicrobial properties, as proposed for some formulations (Elmi et al., 2019), could help reduce antibiotic use. These applications are extremely important in the cattle industry, where sperm cryopreservation and AI are widespread.

Whereas there is ample literature on the use of polyphenols and natural extracts (Ros-Santaella & Pintus, 2021; Santos & Silva, 2020), approaches using agricultural by-products are less studied yet especially promising. These approaches combine the use of natural extracts, free from synthetic substances, while contributing to the circular economy concept and to the revalorization of otherwise waste products. Thus, we explored the application of a blueberry (*Vaccinium corymbosum*) extract obtained from refuse pomace as a supplement to bull semen extenders. Since polyphenols and other substances may be detrimental at excessive concentrations, we conducted a pilot experiment using post-thawed bull semen. Contrary to previous approaches, a “green” methodology yields this extract (Lončarić et al., 2020), with low environmental impact and contributing to more sustainable AI procedures.

## 2 Materials and methods

Semen was collected from eight adult, fertile Holstein bulls using an artificial vagina at the Xenética Fontao breeding center (Lugo, Spain), extended with the same volume of BoviFree extender (Minitüb, Tiefenbach, Germany), cooled to 4 °C, and donated to the researchers. Ejaculates were extended to 92×10^6^/ml in BoviFree, equilibrated, packaged into 0.25-ml straws, and frozen in an IceCube 1800 biofreezer (SY-LAB, Neupurkersdorf, Austria): -5 °C/min to - 10 °C; -40 °C/min to -100 °C; -20 °C/min to -140 °C. Doses were stored in liquid nitrogen, and thawing was performed in a water bath (37 °C, 30 s). The doses were mixed into 3-bull pools, split, and supplemented with 0% (control), 1%, 5%, or 10% aqueous blueberry pomace extract (LIDSA, University of Santiago de Compostela, Spain). The samples in 1.5-ml tubes were incubated at 38 °C up to 5 h.

Motility was assessed hourly with a CASA-mot system (detailed in the supplementary material) using the OpenCASA v. 2 software (Alquézar-Baeta et al., 2019) to obtain total and progressive motility, velocity (VCL, VAP, VSL), linearity (LIN, STR, WOB), ALH, BCF, mean Dance (mDNC), and the fractal dimension.

The experiment consisted of nine pools. Data were analyzed by linear mixed-effects models (R statistical environment v. 4.2), with treatment and incubation time as fixed factors and bull in the random part of the models.

## 3 Results and Discussion

Our results support this blueberry extract as a supplement for bull semen extenders. Figures 1 and 2 summarize each treatment along the incubation time (details in Table S1, supplementary material). After adding 1% of the extract, motility improved both as the proportion of motile cells (Fig. 1a) and in quality (Figures 1b, 1f, 2a, 2b) throughout the incubation time. The fractal dimension (Fig. 2f) is a rarely referenced parameter that reflects the degree to which the sperm trajectory fills a plane and is associated with hyperactivated motility (Boryshpolets et al., 2015), and decreased, indicating lower post-thawing induced hyperactivation (BCF, beat-cross frequency, and mean Dance changed similarly; Figures 2d and 2e). Taken together, these results suggest that supplementation with a low extract amount, possibly due to its polyphenol content, improves post-thawing sperm quality and survivability in cattle.

**FIGURE 1.**
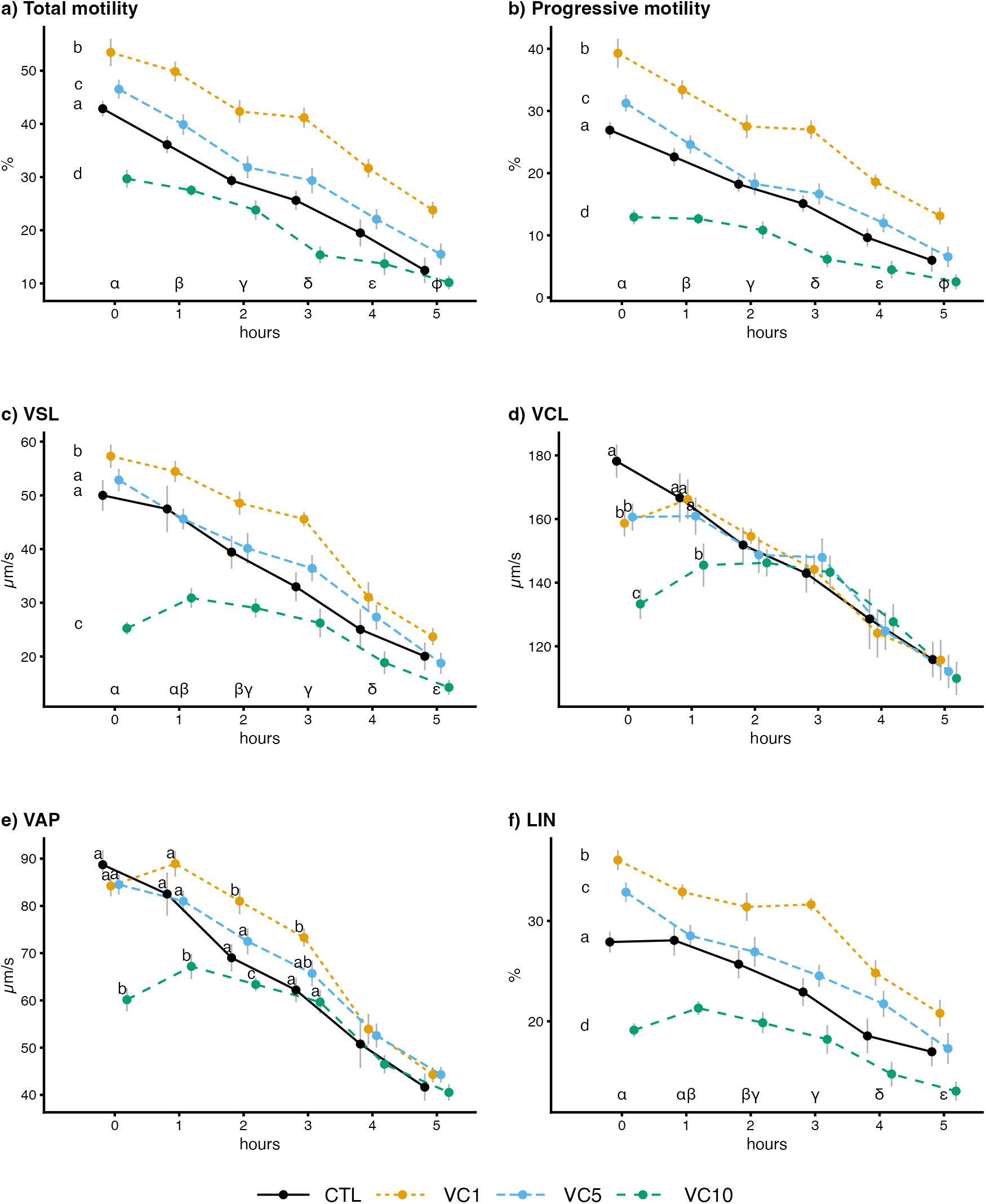
Sperm motility results (mean ± SEM, n = 9). For total motility, progressive motility, VSL, and LIN, the treatment × time interaction was not significant, and the comparison was carried out on main effects (Holms’s adjustment; Latin letters on the left indicate treatment differences and Greek letters indicate time differences). For VCL and VAP, the interaction was significant, with letters indicating differences among treatments at each time point (to prevent cluttering, differences among times are shown in the Suppl. Mat., Table S1). Treatments are CTL: control; VC1: 1% extract; VC5: 5% extract; VC10: 10% extract.

**FIGURE 2.**
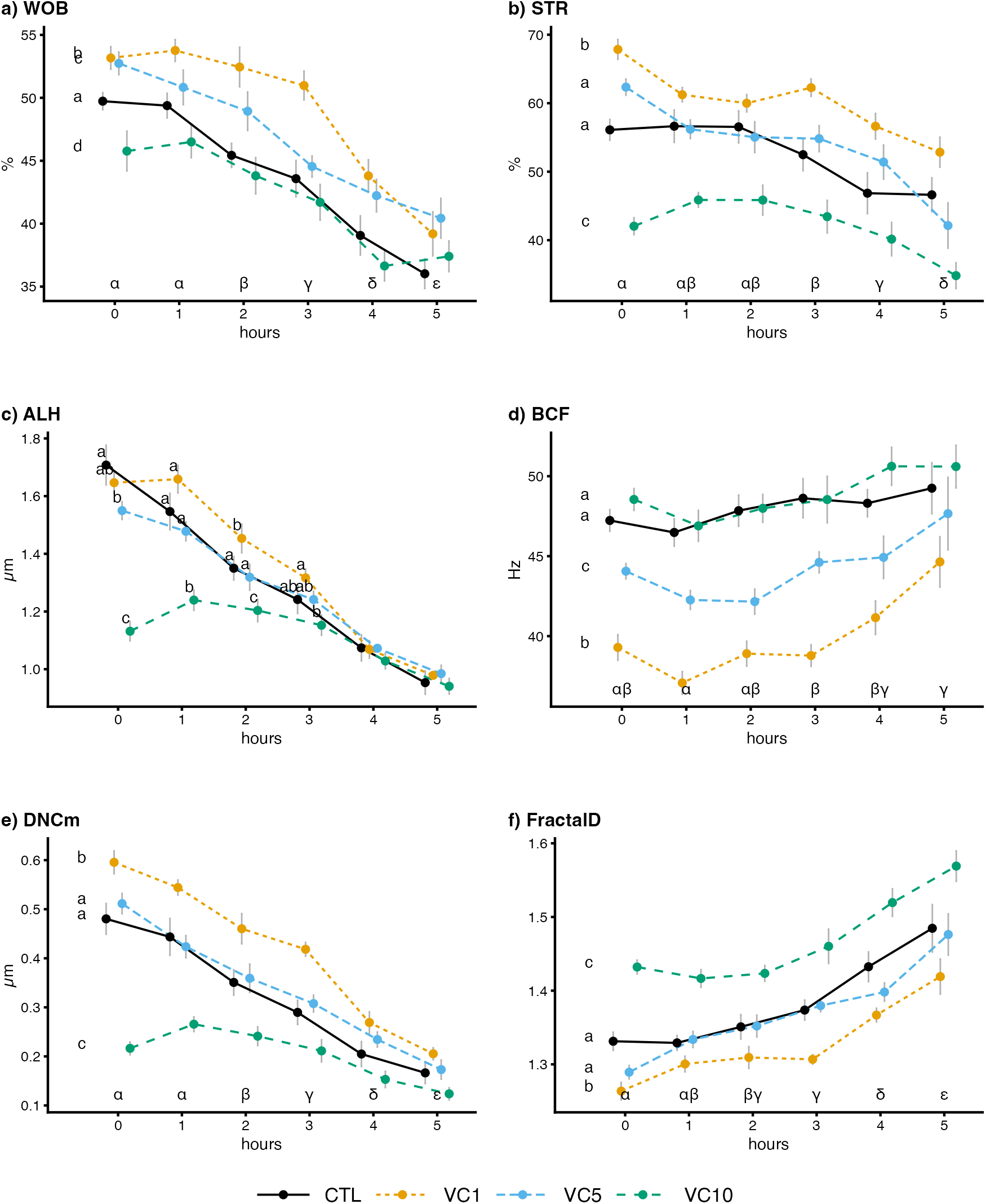
Sperm motility results, see description in Figure 1 (significant interaction only for ALH).

Whereas natural extracts are complex mixtures, which might complicate their use and marketability, this complexity could be an advantage, as shown in applications such as antimicrobials (Gato et al., 2021). Polyphenols and other plant compounds are effective antioxidants, which could explain the protective effects observed here, since oxidative species are major culprits of loss of sperm quality during and after cryopreservation (Sapanidou et al., 2023). Likewise, these natural chemicals are also modulators of membrane pumps and sperm metabolism (Nass-Arden & Breitbart, 1990) and possibly intracellular pathways.

When adding the extract at 5%, we still observed a significant positive effect (P<0.05 in most cases), but with a smaller effect than at 1%. Likely, the complex composition of natural extracts implies a mixture of stimulant and inhibitory effects, whose balance could be less favourable at higher concentrations. Indeed, a physiological level of oxidative species is critical for adequate intracellular signaling (Das & Roychoudhury, 2022), and it could be jeopardized in excessive antioxidative conditions. The motility depression and increased characteristics are compatible with less linear, more hyperactivated motility observed in the 10% treatment, supporting this interpretation.

A major objective of this study was to define safe concentrations for a continuing study of the extract’s applications, since some of the extract’s components or their combinations could exhibit cytotoxicity, and therefore a first definition of a safe range is critical for the development of new formulations. A limitation of this pilot study is that only motility was considered, and therefore, we can only hypothesize molecular effects. Subsequent studies should assess sperm integrity, metabolism, capacitation markers, and chromatin status to elucidate the extract’s effects and better define an optimal concentration.

However, the good results obtained with only 1% offers an excellent prospect for future work, not only for post-thawing improvement but also for refrigerated and cryopreserved storage. Moreover, we must highlight that bull spermatozoa exhibit good tolerance to a relatively high proportion of this extract (between 5% and 10%), which shows promise for exploiting the antimicrobial properties of the blueberry extract (Gato et al., 2021), since these effects are evident at higher polyphenol concentrations.

In conclusion, the blueberry extract tested in this study showed promising protective effects at only 1% in post-thawed bull semen, with no negative effects at 5%. This extract could be useful for enhancing semen extenders in this species, for sperm cryopreservation, and possibly for antibiotic substitution, given its antimicrobial properties. The natural and environmentally respectful origin of this extract makes it attractive in the context of a more sustainable animal production sector.

## Author contributions

GGB designed the experiment, carried out the experiments, and analysed the data; CFH, JFVR, LPM, MVC, and EFA carried out the experiments and analysed the data; BMF, JNC, and JRGM edited the manuscript; ML carried out the experiments and edited the manuscript; FMP coordinated the study, coordinated the data analysis, and drafted the manuscript. All authors read and approved the final manuscript.

## Acknowledgments

The authors thank Xenética Fontao SL (Lugo, Spain) and Indira Álvarez (INDEGSAL, University of León).

## Data Availability Statement

The data that support the findings of this study are openly available in Zenodo at http://doi.org/[doi], reference number [reference number].

## References

Alquézar-Baeta, C., Gimeno-Martos, S., Miguel-Jiménez, S., Santolaria, P., Yániz, J., Palacín, I., Casao, A., Cebrián-Pérez, J. Á., Muiño-Blanco, T., & Pérez-Pé, R. (2019). OpenCASA: A new open-source and scalable tool for sperm quality analysis. PLoS Computational Biology, 15(1), e1006691. 10.1371/journal.pcbi.1006691

Boryshpolets, S., Pérez-Cerezales, S., & Eisenbach, M. (2015). Behavioral mechanism of human sperm in thermotaxis: A role for hyperactivation. Human Reproduction (Oxford, England), 30(4), 884–892. 10.1093/humrep/dev002

Das, A., & Roychoudhury, S. (2022). Reactive Oxygen Species in the Reproductive System: Sources and Physiological Roles. Advances in Experimental Medicine and Biology, 1358, 9–40. 10.1007/978-3-030-89340-8_2

Elmi, A., Prosperi, A., Zannoni, A., Bertocchi, M., Scorpio, D. G., Forni, M., Foni, E., Bacci, M. L., & Ventrella, D. (2019). Antimicrobial capabilities of non-spermicidal concentrations of tea tree (Melaleuca alternifolia) and rosemary (Rosmarinus officinalis) essential oils on the liquid phase of refrigerated swine seminal doses. Research in Veterinary Science, 127, 76–81. 10.1016/j.rvsc.2019.10.014

Gato, E., Perez, A., Rosalowska, A., Celeiro, M., Bou, G., & Lores, M. (2021). Multicomponent Polyphenolic Extracts from Vaccinium corymbosum at Lab and Pilot Scale. Characterization and Effectivity against Nosocomial Pathogens. Plants, 10(12), 2801. 10.3390/plants10122801

Lončarić, A., Celeiro, M., Jozinović, A., Jelinić, J., Kovač, T., Jokić, S., Babić, J., Moslavac, T., Zavadlav, S., & Lores, M. (2020). Green Extraction Methods for Extraction of Polyphenolic Compounds from Blueberry Pomace. Foods (Basel, Switzerland), 9(11), E1521. 10.3390/foods9111521

Nass-Arden, L., & Breitbart, H. (1990). Modulation of mammalian sperm motility by quercetin. Molecular Reproduction and Development, 25(4), 369–373. 10.1002/mrd.1080250410

Ros-Santaella, J. L., & Pintus, E. (2021). Plant Extracts as Alternative Additives for Sperm Preservation. Antioxidants (Basel, Switzerland), 10(5), 772. 10.3390/antiox10050772

Santos, C. S., & Silva, A. R. (2020). Current and alternative trends in antibacterial agents used in mammalian semen technology. Animal Reproduction, 17, e20190111. 10.21451/1984-3143-AR2019-0111

Sapanidou, V., Tsantarliotou, M. P., & Lavrentiadou, S. N. (2023). A review of the use of antioxidants in bovine sperm preparation protocols. Animal Reproduction Science, 251, 107215. 10.1016/j.anireprosci.2023.107215

Schulze, M., Nitsche-Melkus, E., Hensel, B., Jung, M., & Jakop, U. (2020). Antibiotics and their alternatives in Artificial Breeding in livestock. Animal Reproduction Science, 220, 106284. 10.1016/j.anireprosci.2020.106284

