## Supplemental methods for "A low concentration of a sustainably obtained blueberry extract improves the post-thawing motility of cryopreserved bull spermatozoa"

### CASA-mot procedures and configuration

The CASA-mot (Computer Assisted Analysis System for motility) consisted of a Nikon E600 microscope (Nikon, Tokyo, Japan) with a warmed stage at 38 °C, with an acA1920-155uc camera (Basler, Ahrensburg, Germany) attached (no intermediate magnification). Sample and video analyses followed routine protocols for bull spermatozoa in our laboratory. Sperm samples were loaded in a modified Makler counting chamber (25 µm height), and images were acquired at ×10 magnification (negative phase contrast). The Pylon v. 5 software (Basler) controlled image acquisition, with parameters displayed in Table S2. Images were stored as uncompressed AVI files for further analysis, using a hierarchical directory structure that included the pool, sampling time, and treatment for each sample in the directories' names. At least three fields per sample were acquired.

The AVI videos were processed with the OpenCASA v. 2 software (Alquézar-Baeta et al., 2019), available at https://github.com/calquezar/OpenCASA, as an ImageJ v. 1.54g plugin (https://imagej.net/), using the multiple directory option and the configuration shown in Table S3. Individual and summary motility data were stored in CSV files, which were further processed using the OpenCASA R package (Martínez-Pastor, 2026), available in https://codeberg.org/fmartinezpastor/OpenCASA/.

OpenCASA, ImageJ, the OpenCASA R package, and R are free and open source software (FOSS), with code available at the cited URL.

The OpenCASA2tables() function in the OpenCASA R package was used with the default parameters, except for cond_prog, which was set to vap = 15 and str = 50. The function produced, for each sample, the parameters described in Table S4, which were subsequently used in the statistical analysis with linear mixed-effects models. This dataset is available at http://doi.org/[doi].

Table S2. Configuration of the Basler Pylon software for acquiring bull sperm motility videos.

| Parameter | Value |
|---|---|
| Output format | AVI |
| Use compression | No |
| Record a frame every | 1 frames |
| Stop recording after | 1200 miliseconds |
| Width | 800 pixels |
| Height | 800 pixels |
| Pixel format | Mono 8 |
| Exposure time | 180.0 |
| Exposure auto | Off |
| Acquisition frame rate | 240 |

Table S3. Configuration of the OpenCASA v. 2 software for analyzing the bull sperm motility AVI videos.

| Parameter | Value |
|---|---|
| Microns per Pixel | 588 |
| Minimum cell size ($\mu m^2$) | 15.0 |
| Maximum cell size ($\mu m^2$) | 120.0 |
| Progressive motility (STR>%) | 50.0 |
| Progressive motility (VAP>) | 15.0 |
| Minimum VCL (µm/s) | 10.0 |
| VCL lower threshold (µm/s) | 40.0 |
| VCL upper threshold (µm/s) | 50.0 |
| Frame Rate (frames/s) | 200.0 |
| Minimum Track Length (frames) | 15 |
| Maximum displacement between frames (µm) | 6.0 |
| Window Size (frames) | 9 |
| Print XY coords | No |
| Save Video Output | Yes |

Table S4. Description of the CASA parameters (definitions taken from Boyers et al., 1989; fractal dimension from Alquézar-Baeta et al., 2019, and Mortimer et al., 1996).

| Parameter | Definition |
|---|---|
| Total motility | Percentage of spermatozoa exceeding a minimum VCL threshold |
| Progressive motility | Percentage of spermatozoa exceeding a minimum VAP and STR threshold |
| VCL | Curvilinear velocity: the time-average velocity of the sperm head along its actual trajectory |
| VSL | Straight-line velocity: the time-average velocity of the sperm head along a straight line from its first detected position to its last detected position |
| VAP | Average-path velocity: the time-average velocity of the sperm head along its average trajectory, defined as the mathematical smoothing of the actual trajectory |
| LIN | Linearity: the linearity of the curvilinear trajectory (VSL/VCL) |
| WOB | Wobble: the degree of oscillation of the actual sperm trajectory around the average path (VAP/VCL) |
| STR | Straightness: the straightness of the average path (VSL/VAP) |
| ALH | Amplitude of the lateral head displacement: the amplitude of variations of the actual sperm-head trajectory along its average trajectory (OpenCASA provides mean and maximum ALH values; mean ALH is reported in the present work) |
| BCF | Beat-cross frequency: the time-average rate at which the actual sperm trajectory crosses its average path trajectory; an extrapolation of the flagellar beat frequency |
| DNCm | DanceMean: a measure of the pattern of sperm motion (ALH/LIN) |
| Fractal dimension | Degree to which a line fills a plane; a measurement of the regularity of the trajectory, with a low value showing regularity and a high value showing randomness (potentially related to hyperactivation) |

### References


Alquézar-Baeta, C., Gimeno-Martos, S., Miguel-Jiménez, S., Santolaria, P., Yániz, J., Palacín, I., Casao, A., Cebrián-Pérez, J. Á., Muiño-Blanco, T., & Pérez-Pé, R. (2019). OpenCASA: A new open-source and scalable tool for sperm quality analysis. PLoS Computational Biology, 15(1), e1006691. https://doi.org/10.1371/journal.pcbi.1006691

Boyers, S. P., Davis, R., & Katz, D. F. (1989). Automated semen analysis. Curr. Probl. Obstet. Gynecol. Fertil., 12(5), 165–200.

Martínez Pastor, F. (2026). OpenCASA: Compiling OpenCASA Results Files into Annotated Tables. R package version 1.0.0, https://codeberg.org/fmartinezpastor/OpenCASA/

Mortimer, S. T., Swan, M. A., & Mortimer, D. (1996). Fractal analysis of capacitating human spermatozoa. Human Reproduction, 11(5), 1049–1054. https://doi.org/10.1093/oxfordjournals.humrep.a019295
